# Diverse Intracellular Trafficking of Insulin Analogs by Machine Learning-based Colocalization and Diffusion Analysis

**DOI:** 10.1101/2024.12.12.628100

**Authors:** Sara Vogt Bleshøy, Jacob Kæstel-Hansen, Annette Juma Nielsen, Narendra Kumar Mishra, Knud J. Jensen, Nikos S. Hatzakis

**Author notes:** These authors contributed equally (Sara Vogt Bleshøy, Jacob Kæstel-Hansen).

## Abstract

Insulin signaling is crucial for maintaining cellular function and systemic homeostasis, with its dysregulation leading to metabolic disorders in particular diabetes. While insulin analogs are essential in type 1 diabetes treatment, their intracellular trafficking and sorting compared to endogenous insulin remains largely unresolved at the molecular level. Understanding these are important in improving therapeutics and guiding future drug development. However, current methods rely on static imaging and bulk receptor assays conducted in non-physiological conditions, which often disrupt native insulin signaling and provide only snapshots that fail to capture the temporal dynamics of insulin trafficking and characteristic sorting pathways. Here, we directly observe insulin trafficking and intracellular sorting and compare the characteristics of Atto655 labeled recombinant human insulin (HI^655^) with insulin aspart (IAsp^655^), a clinically approved, rapid-acting insulin analog. We developed a combined approach integrating Colocalization Fingerprinting, a machine learning framework for reliable, time-resolved colocalization analysis, together with our recently developed deep learning-assisted single-particle diffusional analysis (DeepSPT). Our analysis revealed subtle, yet significant differences in intracellular behavior between IAsp^655^ and HI^655^, particularly in diffusional behavior and lysosomal colocalization, highlighting the potential of our approach to decipher subtle differences in intracellular trafficking and sorting characteristics. In addition to contributing to a more detailed understanding of biology of insulin analogs and intracellular sorting, we provide a reliable machine-learning methodology to study intricate cellular processes.

## Introduction

Insulin signaling is a complex process, where production, secretion, receptor binding, intracellular trafficking, and degradation are meticulously regulated to ensure proper glycaemic response, cell growth, and metabolic and mitogenic homeostasis.^1,2^ Dysregulation of this pathway triggers severe metabolic diseases in particular diabetes mellitus, and with global rise in diabetes cases, pharmaceutical insulin analogs are increasingly important.^3–5^ Although all pharmaceutically available analogs are well described in clinical trials, studies quantifying their internalization, intracellular trafficking and sorting - beyond receptor binding kinetics, bulk uptake, and degradation assays in fixed or unresponsive cells,^6–10^ are limited. Even minor changes in the amino acid sequence of the analog may alter the receptor binding affinity, association, and disassociation, impacting the internalization and intracellular sorting.^6,7,11^ Therefore, understanding these trafficking mechanisms are crucial to gain insights into intracellular signaling of insulin and its analogs, and to inform the design of pharmaceuticals for prolonged, improved treatments.^12–14^

Current understanding of intracellular behavior of insulin and its analogs is predominately based on snapshots obtained through fluorescence and electron microscopy combined with qualitative^15–19^ or pixel-based analysis,^20,21^ or inferred from bulk receptor binding assays.^15,17,22^ These approaches often rely on non-physiological conditions such as purified receptors, fixed or serum-starved cells, pH controlled environments, or cold conditions that disrupts endocytosis and native insulin signaling.^21,23,24^ As a result, temporal dynamics intrinsic to internalization and intracellular sorting are often masked. Live-cell fluorescence microscopy and single particle tracking (SPT) of insulin variants and cellular organelles offer methodologies to capture spatiotemporal dynamics of insulin trafficking pathways and diverse intracellular colocalization partners under physiological conditions.^24–26^ While labeling protein and cellular compartments for intracellular colocalization assays is well-established^27–30^, identifying genuine colocalization partners remains challenging due to incidental proximity generated in densely packed, compartmentalized cellular environments^31–33^ and by anomalous diffusion, which creates large spatiotemporal variability and periods of low mobility, prolonging incidental proximity.^34,35^ Furthermore, technical artifacts, such as chromatic offset and localization errors obscure colocalization by altering the perceived proximity of entities.^36–39^

Several approaches to colocalization analysis tackling the above challenges exist, ranging from visual inspection to statistical analysis.^20,27,28,40–53^ However, colocalization is most often condensed to a single value, thus also masking particle heterogeneity and temporal variations important for deciphering intracellular function. A few approaches for time-resolved colocalization exist, among them Deschout et al.^54^ and Dupont et al.^55^ However, the methods remain sensitive to crowded cellular environments, localization errors, and chromatic offsets.^54,55^

Here, we provide real-time spatiotemporal observation of insulin internalization and recycling to compare human insulin (HI) and insulin aspart (IAsp), a rapid acting variant, which differs from HI by a single point mutation from proline to aspartic acid on the B-chain (B28Asp). IAsp (commercially NovoLog or NovoRapid) displays a faster onset of action and plasma clearance rates, but similar function and receptor-binding properties to recombinant HI.^10,11,56,57^ We use SPT to track in real-time the spatiotemporal localization of Atto655 labeled Human Insulin (HI^655^) and labeled Insulin Aspart (IAsp^655^) in parallel with fluorescently labeled endolysosomal and recycling compartments. To overcome the challenges of analysis of intracellular diffusion and time-resolved colocalization, we developed and utilized two machine learning frameworks: DeepSPT (Kæstel-Hansen et al.^58^), a deep learning method for extracting biological insights from heterogeneous single-particle diffusion, and Colocalization Fingerprinting, a machine learning-driven methodology designed for time-resolved differentiation between incidental and meaningful biological colocalization. The combined machine-learning approach identified intracellular diffusional differences between the two investigated insulin analogs and revealed subtle yet significant differences in intracellular sorting between IAsp^655^ and HI^655^. Importantly, we identify an increased localization of HI^655^ with lysosomes in HEK293 cells, but find no significant differences in the sorting of the variants in HeLa cells, highlighting the potential of our approach to decipher subtle differences in intracellular trafficking and sorting characteristics. This approach establishes a robust, open-source framework to examine dynamic and intricate intracellular interactions in complex cellular environments for physiological and pathological conditions.

## Results

### Probing intracellular motion by machine learning

To investigate intracellular trafficking differences between human insulin and insulin aspart, we analyzed their diffusion and colocalization with post-endocytic markers in HEK293 cells. Separate populations of cells were fluorescently labeled by either Rab5a-emGFP (early endosomes), Rab7a-emGFP (late endosomes), LysoTracker Green (lysosomes/endolysosomes), or transferrin (probing both slow and fast recycling pathways^59–61)^ together with an Atto655 labeled insulin variant (Fig. 1a and b). For each condition, one cellular compartment was tagged and one insulin analog added, giving 8 total conditions (2 insulin variants, 4 cellular markers) to follow intracellular behavior in real time. Combined, we studied ∼56500 insulin and ∼255000 endosomal trajectories, totaling ∼8.75 and ∼26.7 million localizations, respectively.

**Figure 1.**
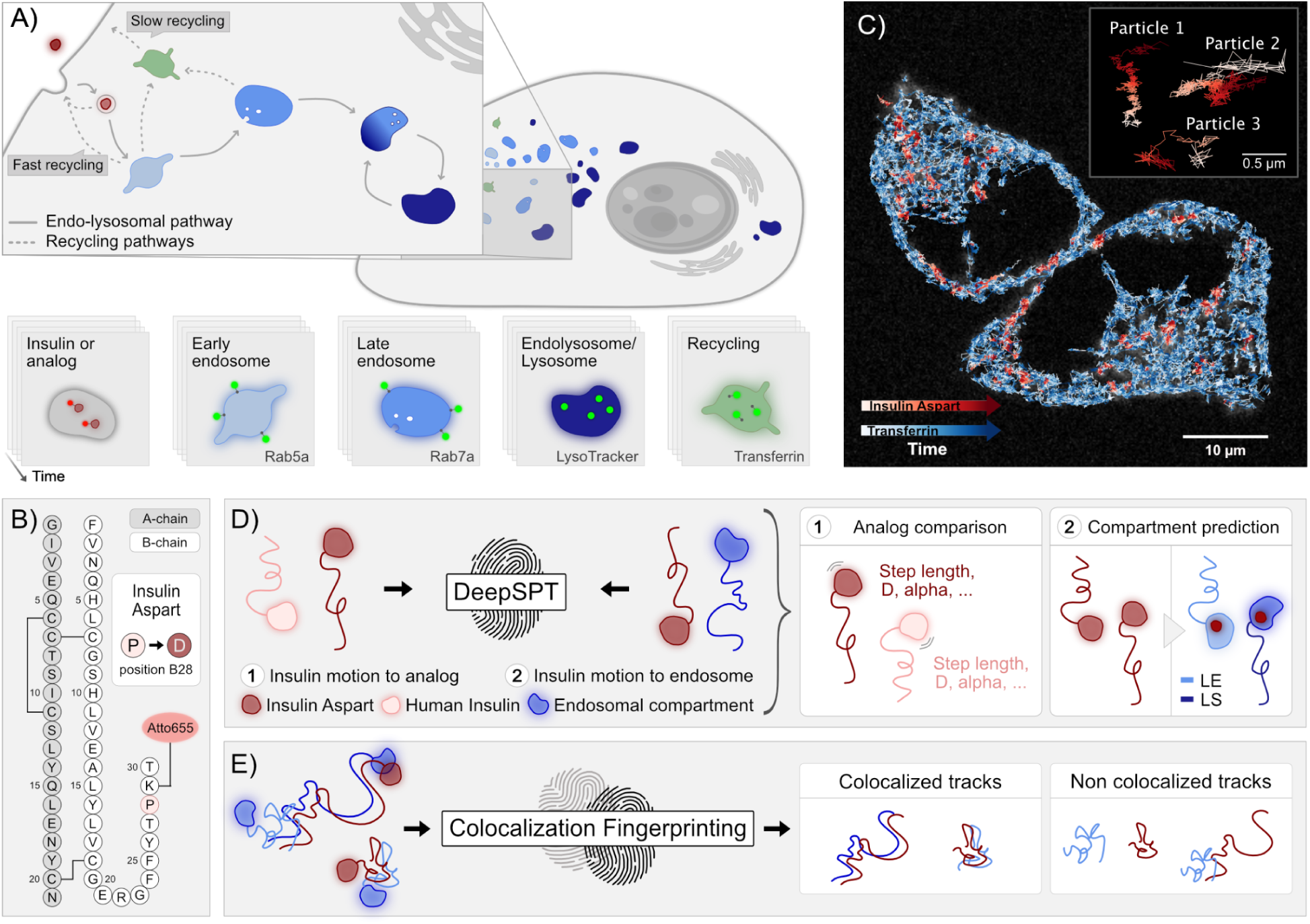
Machine learning framework for live-cell single-particle tracking elucidates variations in intracellular trafficking of insulin analogs. **A)** A schematic representation of the live-cell assay used to investigate intracellular trafficking patterns. Cells are incubated with labeled insulin (see primary structure in **B**) and a fluorescent marker probing an endosomal compartment; early endosomes by Rab5a, late endosomes by Rab7a, endolysosomes and lysosomes with LysoTracker or transferrin to probe recycling pathways. Temporal 2D molecular movies in two channels are collected to compare insulin behavior with the given endolysosomal or recycling compartment. **B)** Primary structure of Atto655 labeled Human Insulin (Atto655 and insulin are not to scale) highlighting the single point mutation from proline to aspartic acid on position B28, of the fast-acting insulin analog, insulin aspart. **C)** Representative Spinning disk confocal microscope image of dual-labeled HEK293 cells (scale bar 10 µm). Live-cell imaging produces a set of x, y, and t, localizations for each particle, yielding a dataset of single particle trajectories (inset, scale bar 0.5 µm). Here, cells are overlaid with the corresponding single particle trajectories of insulin aspart (red, N=269) and transferrin (blue, N=2911). **D)** Trajectories are fed to DeepSPT, a deep-learning framework, which quantifies diffusional variation between entities, here the two insulin variants (1), and classifies them according to the most similar endolysosomal diffusion (2). LE = Late endosomes, LS = Lysosomes. **E)** Colocalization Fingerprinting, a machine learning pipeline for reliable and automatic identification of colocalization based on temporal consistent proximity, and spatial and diffusional features.

Quantification of colocalization is not trivial as it is challenged by crowded cellular environments (Fig. 1c), chromatic aberrations (Supplementary Fig. 6) and localization errors, which cause coincidental detections and limit existing pixel-based approaches.^62,63^ We propose to evaluate colocalization by combining the spatiotemporal development of particles, with diffusional analysis and intracellular proximity. Building on previous work from our group, we utilize DeepSPT (Kæstel-Hansen et al.^58^), a deep learning-based toolbox to rapidly, accurately, and automatically analyze heterogenous single-particle tracking diffusion to deconvolute diffusional variations between the two insulin species (Fig. 1d). Additionally, we introduce Colocalization Fingerprinting as a new improved approach to study colocalization (Fig. 1e). Together, they form a synergistic framework to elucidate variations in intracellular interactions,internalization, sorting, and degradation pathways, to quantitatively describe differences in intracellular behavior between the two insulin variants.

### Slight Variations in Intracellular Diffusion Between Insulin Analogs

To compare the intracellular behavior of the two insulin variants, we analyzed their intracellular motion by examining the displacement of each particle between consecutive frames (Fig. 2b). IAsp^655^ displays statistically significant longer step lengths as determined by a one-sided Welch’s t-test (see Supplementary Fig 7 and Supplementary Table 2) and thereby faster motion, with median average step lengths of 0.18 µm as compared to 0.13 µm for HI^655^. Given the only difference between the insulin variants is a single point mutation (Fig. 1b) this significant difference in intracellular diffusional behavior is interesting.

**Figure 2.**
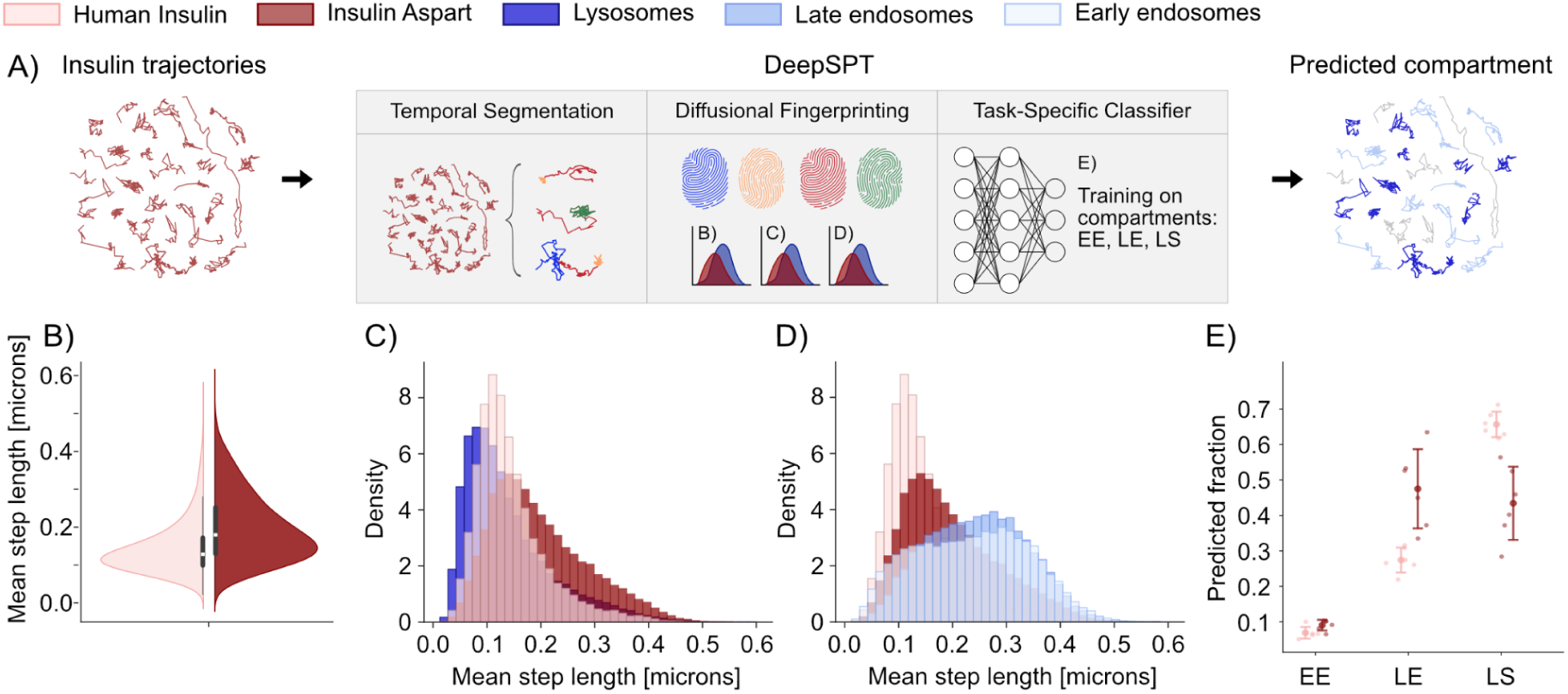
Insulin analogs display diverse intracellular diffusional behavior. **A)** Schematic illustration of the DeepSPT pipeline for descriptive featurization of trajectories and diffusion-based identity prediction. Firstly, trajectories are fed to DeepSPT, which, for each trajectory, temporally segments diffusional behavior and converts the segmented trajectories into an extensive feature representation, termed a diffusional fingerprint. Subsequently, this diffusional fingerprint informs a classifier, trained to differentiate endosomal compartments, in categorizing trajectories to a specific compartment identity. **B)** Trajectory mean steplength analysis for HI^655^ (pink) and IAsp^655^ (red). The trajectories are the sum from 6 biological replicates per insulin variant with medians 0.13 µm and 0.18 µm for HI^655^ and IAsp^655^, respectively (See Supplementary Table 2 and Supplementary Fig. 7 for individual biological replicates). **C)** Histograms of mean step lengths of trajectories above 20 time points are shown for the insulin analogs with lysosomes (LysoTracker), and **D)** endosomes (Rab5a, light blue and Rab7a, blue). Number of trajectories: N_HI_ = 20901, N_IAsp_ = 35602, N_EE_ = 45351, N_LE_ = 23072, N_LS_ = 55174. **E)**. Fraction of tracks of either insulin analog, predicted by DeepSPT to diffuse like the respective compartment for trajectories above 50 time points. Errors depict standard deviation across technical replicates. EE = Early endosomes, LE = Late endosomes, LS = Lysosomes. Number of trajectories: N_HI_ = 11925, N_IAsp_ = 17457, N_EE_ = 18980, N_LE_ = 17725, N_LS_ = 13859.

We compared the insulin step lengths to the ones of the compartments in the endolysosomal pathway, to link variations in step lengths to cellular localization. In Fig. 2c and 2d, the mean step length distributions of both insulin analogs are compared to those of lysosomes (Fig. 2c), and early and late endosomes (Fig. 2d). Lysosomes display shorter displacements than the two remaining compartments, suggesting HI^655^ motion to be more like the slower moving compartments such as the lysosomes, and IAsp^655^ to behave more like the faster endocytic compartments earlier in the pathway, indicating differences in cellular localization.

To explore this further, we employed DeepSPT^58^, which based on diffusional features predict intracellular localization of particles. The model was trained on the diffusional patterns of the three labeled compartments; early endosomes, late endosomes, and lysosomes and evaluated on their diffusional similarity to the insulin trajectories (Fig. 2a). Consistent with the displacement analysis in Fig. 2b-d, IAsp^655^ is predicted to diffuse similarly to late endosomes while HI^655^ diffused more similarly to lysosomes (Fig. 2e). Neither of the insulin variants were predicted to behave like early endosomes (Fig. 2e). This could either be from receptor saturation limiting new internalization events after longer incubation times, or due to the diffusional similarity of early and late endosomes, challenging their differentiation and favoring prediction of the late endosome class, which is more distinct from lysosomes (Supplementary Fig. 8).

The direct recording of diffusional behaviors revealed subtle but significantly different intracellular sorting of the insulin variants. This is surprising, as studies so far have shown IAsp to resemble HI in affinity and binding constants with the two main binding partners of insulin action; the insulin receptor and the IGF-1 receptor.^7,8^ While only a small difference in the insulin receptor off-rate between the analogs has been reported, which was deemed inconsequential for native-like function^10,64^, our findings suggests differences in intracellular motion and lysosomal colocalization, arriving from their structural differences.

### Accurate Differentiation of Genuine and False Colocalization by Colocalization Fingerprinting

Intracellular colocalization analysis is a daunting challenge due to the densely crowded environment, heterogeneous diffusion, and technical artifacts that obscure coincidental proximity of non-interacting particles from true interactions.^31–33,37,38^ To tackle these challenges we introduce Colocalization Fingerprinting, a machine learning-based framework, which identifies colocalizing partners based on their similarity in proximity and motion and provide time-resolved colocalization annotation. The framework employs three sequential modules: The first module (M1) proposes potential colocalizing pairs of trajectories based on temporally consistent proximity, thereby temporarily evaluating colocalization while ensuring spatial adjacency and minimizing the inclusion of spurious proximities (Fig. 3a, see Methods). The second module (M2) transforms each identified trajectory pair into a comprehensive similarity representation, termed a colocalization fingerprint. This 23-dimensional fingerprint is derived from spatial characteristics such as inter-trajectory distance statistics, trajectory similarity metrics, spatial coordinate correlation, and diffusional features including step statistics, trajectory geometry, and temporal diffusional behavior provided by DeepSPT (Fig. 3a, see Methods).^58^ The third module (M3) employs a machine learning classifier leveraging the colocalization fingerprint of each trajectory pair to assess the authenticity of each proposed colocalization pair. Additionally, a set of user-defined, fail-safe filters is implemented to further improve reliability (Fig. 3a, see Methods).

**Figure 3.**
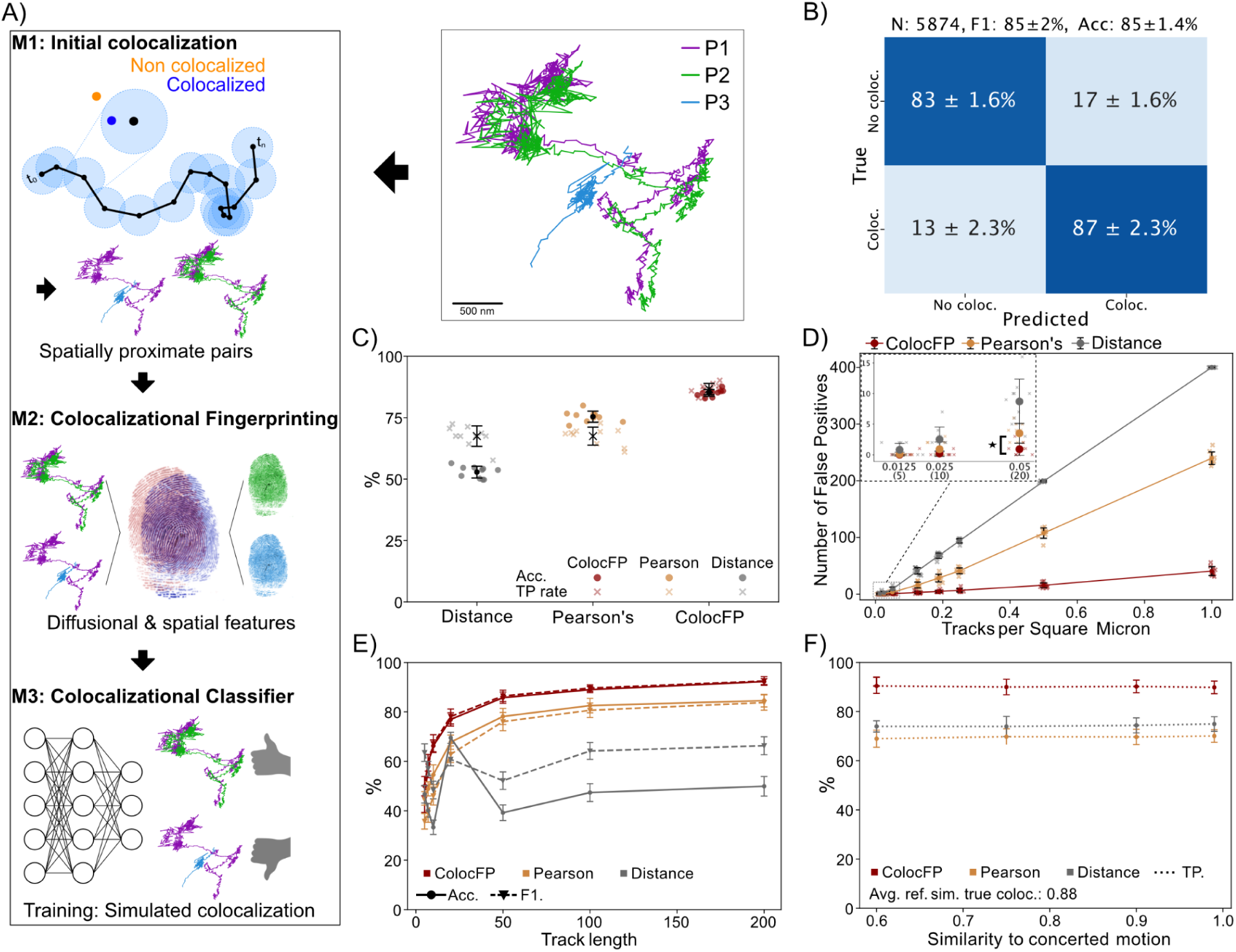
Colocalization Fingerprinting accurately differentiates true and false colocalization in crowded environments. **A)** Schematic representations of three particles (Top center), P1, P2, and P3, are shown. P1 and P2 (purple and green) illustrate concerted motion and genuine colocalization, while P1 and P3 (purple and blue) display incidental proximity, thus false colocalization. Top left, depicts the Colocalization Fingerprinting pipeline: M1 identifies pairs of trajectory segments that colocalize based on persistent proximity in time; M2 constructs the colocalization fingerprint of each pair, utilizing diffusion and spatial characteristics; M3 consists of a machine learning classifier and fail-safe filters to distinguish between true and false colocalization. **B)** The confusion matrix for M3 in differentiating simulated true (Coloc.) and false (No coloc.) colocalization (see Methods) as assessed in a ten-fold cross-validation. Accuracy (Acc.), number of tracks (N), and F1-score (F1) provide additional classification metrics. Errors depict standard deviation across the ten folds. **C)** The accuracy and true positive rate (TP rate) for Colocalization fingerprinting (ColocFP) and benchmark approaches in a ten-fold cross-validation. Pearson’s represents utilizing a cutoff at 0.5 in Pearson’s correlation.^49^ Distance denotes solely relying on the average inter-trajectory distance to mimic a search-range-based approach. Errors depict standard deviation across the ten folds. **D)** Evaluation of Colocalization Fingerprinting and benchmark approaches in filtering out false colocalization in an enclosed space (20×20 μm) with an increasing density of non-interacting particles (see Methods, Supplementary Fig. 4). The number of identified colocalizing segments directly denotes the false positive rate and the susceptibility to crowding. Zoom-in depicts the low-density regime for visual clarification. Colocalization Fingerprinting consistently identifies significantly fewer false positives down to densities of 0.05 tracks/μm^2^ (20 tracks per 400 μm^2^), as evaluated by a two-sided Welch’s t-test (p-value<0.0007, N=10 per approach). In comparison, experimentally observed densities of endolysosomal and recycling compartments were identified to range from 0.03-0.15 tracks/μm^2^, with a mean ± sem of 0.101 ± 0.006 tracks/μm2 (see Methods). The diffusion coefficients, track durations, and search range used to construct the simulations are provided in Methods. Errors depict standard deviation across ten independent crowding simulations. **E)** Classification performance of Colocalization Fingerprinting and benchmark approaches for various track lengths. Colocalization Fingerprinting displays significantly greater accuracy across all trajectory durations by quantification using a one-sided Welch’s t-test (all p-values<=0.034, N=10 per approach). Errors depict standard deviation across the ten folds. **F)** Accuracies for Colocalization Fingerprinting and benchmark approaches in conditions where false colocalization increasingly resembles perfectly concerted motion, thus increasingly obscuring the differences between true and false colocalization. Errors depict standard deviation across the ten folds.

To validate the effectiveness and reliability of Colocalization Fingerprinting in evaluating colocalization, we employ four evaluation schemes based on simulations as compared to benchmark approaches.

Firstly, we quantify Colocalization Fingerprinting’s classification accuracy on simulated, heterogeneously diffusing, truly colocalized trajectories perturbed by technical artifacts as well as distinct, independent trajectories representing spurious proximity of non-interacting particles (see Methods). Most confusion between classes with our method stems from short trajectories (see Supplementary Fig. 9) and the main source of its improvement over the Pearson’s correlation benchmark is an increased true positive rate (see Supplementary Fig. 10 for additional benchmarking of classification). Proximity alone results in a random baseline classifier accuracy of 53 ±2% while Pearson’s correlation reaches 75 ±2%. In contrast, Colocalization Fingerprinting obtains an accuracy of 85 ±1.4%, significantly outperforming the competition (Fig. 3b,c).

Secondly, we assess the reliability of Colocalization Fingerprinting in densely populated environments by quantifying the number of colocalization pairs identified upon increasing density of heterogeneously diffusing, non-interacting particles (see Methods, Supplementary Fig. 4). In this analysis, Colocalization Fingerprinting significantly decreases the incidences of false positive detections compared to a search range-based strategy and Pearson’s correlation (Fig. 3d) down to sparsely populated conditions of 0.05 tracks/μm^2^ (see Supplementary Fig. 5), as verified by a two-sided Welch’s t-test. In comparison, the observed densities of endolysosomal and recycling compartments in experiments were identified to range from 0.03-0.15 tracks/μm^2^ (mean ± sem of 0.101 ± 0.006 track/μm^2^, see Methods), yet higher densities for smaller objects is highly likely. The improvement of Colocalization Fingerprinting is particularly pronounced in highly crowded settings (1 track/μm^2^), where it reduces false positive detections by up to approximately tenfold, highlighting its reliability in crowded environments.

Thirdly, we evaluated the sensitivity of Colocalization Fingerprinting to the duration of proposed colocalizing segments by quantifying classification accuracy across various durations (see Methods). Colocalization Fingerprinting demonstrates accuracies of 59 ±3% at 7 time points, 66 ±2% at 10 time points, and up to 92 ±1.2% at 200 time points, while coordinate Pearson’s correlation obtains 49 ±3%, 55 ±4%, and 85 ±3%, and distance-based evaluation 47 ±4%, 40 ±3%, and 50 ±4% for the same number for time points (Fig. 3e). Colocalization Fingerprinting is the only approach to achieve non-random accuracies at just 7 time points and consistently outperforms the benchmarks across all trajectory durations by quantification using a one-sided Welch’s t-test.

Lastly, we investigate the sensitivity of Colocalization Fingerprinting to trajectories of non-interacting particles that progressively exhibit diffusional properties more akin to perfectly concerted motion (see Methods). As false colocalization begins resembling perfectly concerted motion the important metric becomes true positive rate, as the false positive rate will undoubtedly increase and the accuracy may drop, it remains important to retain the true positives. The stress test will eventually lead to all pairs being predicted as colocalizing, but this is not considered undesirable behavior based on the assumption that in realistic scenarios, non-interacting particles do not exhibit more concerted motion than interacting particles. In this scenario, Colocalization Fingerprinting maintains the high true positive rate (see Fig. 3f) and also consistently outperforms benchmark methods.

Combined, these results underline that solely relying on distance is unreliable in differentiating true colocalization from incidental proximity. Increasing a search range with Pearson’s correlation improves the reliability, yet remains challenged especially in crowded environments. In contrast, Colocalization Fingerprinting utilizing spatiotemporal characteristics consistently outperforms the benchmark approaches and provides reliable and robust colocalization.

### Insulin Analogs Exhibit Varying Propensities for Endosomal Colocalization in HEK293

Next, we focused on the dual-color live cell SPT to characterize the endosomal colocalization of the two insulin variants (Fig. 3). From the experimental data, it is evident why proximity alone often provides coincident detection of non-interacting entities due to crowding and anomalous diffusion, convolved by imaging artifacts (see Fig. 1c and Supplementary Fig. 6). The example in Fig. 4a (and additional examples in Supplementary Fig. 11,12) shows a trajectory pair, which remains constantly within ∼500 nm throughout its lifetime, resulting in a falsely classified colocalizing pair with classical methods, even though its diffusional and spatial characteristics are different (see Supplementary Fig. 6). Fig. 4b displays a representative example of true colocalization. The particles clearly display similar diffusional and spatial characteristics, albeit obscured by localization errors and a chromatic offset producing an inter-particle distance similar to the false colocalization example in Fig. 4a (∼400-500 nm). Colocalization Fingerprinting reliably differentiates incidental proximity from biologically relevant interactions in these experimental examples (Fig. 4a,b, and more examples Supplementary Fig. 11,12) predicting only Fig. 4b, and not Fig. 4a, as true colocalization. Importantly, the examples support the notion that true colocalization exhibits concerted motion, and while noise and spurious proximity may obscure the differentiation for metrics as Pearson’s correlation (Supplementary Fig. 12), Colocalization Fingerprinting reliably differentiates false and true colocalization in complex intracellular environments.

**Figure 4.**
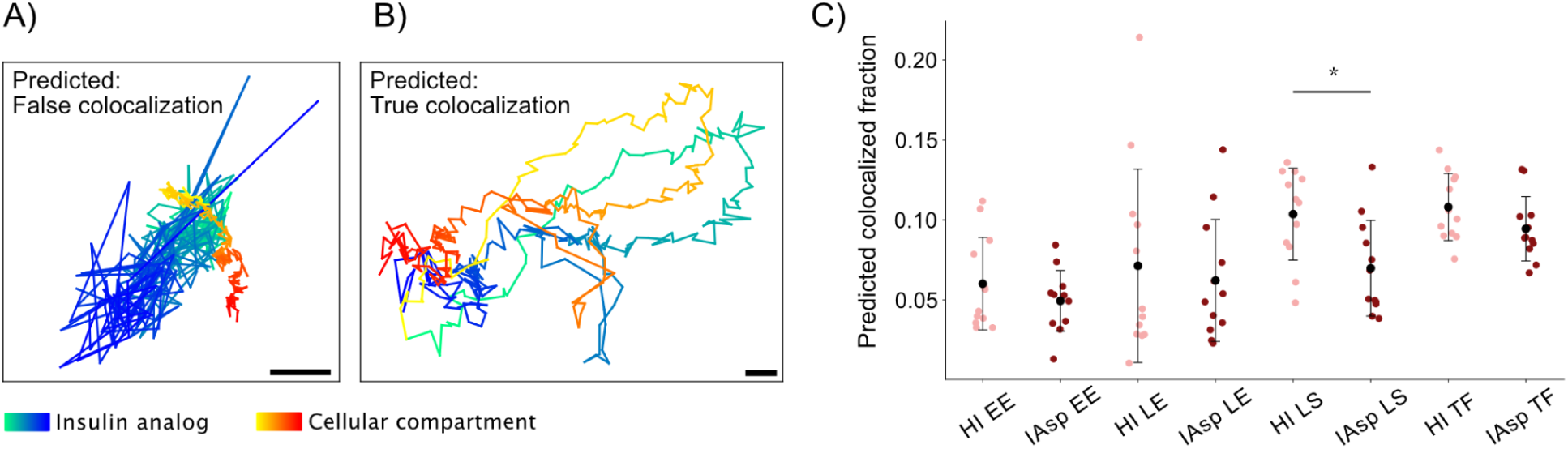
Human insulin and the fast-acting analog, insulin aspart, exhibit varying intracellular colocalization propensities. **A)** Proposed colocalizing trajectory pair from dual-color SPT of insulin and endosomal compartments (see Methods). Proposals are based on a specified search range and the associated Colocalization Fingerprinting prediction is shown in the plot (see Methods). This example showcases how crowded environments lead to spatially proximate trajectories that do not display concerted motion or indications of colocalization. **B)** Proposed colocalizing pair as in (**A**) from dual-color SPT based on a specified search range together with its classification by Colocalization Fingerprinting. **A-B)** Additional examples in Supplementary Fig. 11 and 12. Trajectories are colored by increasing time from lighter to darker tones. Scale bar is 200 nm. **C)** Fraction of insulin tracks colocalizing with a given endosomal compartment observed across eight distinct two-color SPT experiments (see Methods). Colocalization is predicted and reported by Colocalization Fingerprinting with each point denoting a technical replicate (12 technical replicates across 2 biological replicates). HI = human insulin, IAsp = Insulin aspart, EE = Early endosomes, LE = Late endosomes, LS = Lysosomes, TF = Transferrin. The two insulin variants’ lysosomal colocalization propensity differs significantly, as evaluated by a two-sided Welch’s t-test (p-value < 0.01, N=12 per insulin variant). Error bars depict standard deviation across technical replicates.

Quantification of endosomal colocalization of the two insulin analogs across 12 dual-color single-particle tracking experiments per condition (Fig. 4c, two biological replicates, see Methods), revealed a slight, yet significant difference in lysosomal colocalization between the two insulin analogs (two-sided Welch’s t-test). This supports the finding from DeepSPT (Fig. 2e) and suggests differences in intracellular sorting, trafficking and possibly degradation characteristics between the two insulin variants.

Previous studies report slightly lower receptor dissociation constant (K_d_) of IAsp as compared to HI, with the K_d_ of IAsp 81 ±8% of the K_d_ of HI,^10,64^ which appears to correlate with the lower lysosomal colocalization for IAsp^655^. This characteristic could prompt IAsp to follow the recycling pathway of the insulin receptor more frequently, rather than releasing and continuing through the endolysosomal machinery. While transferrin was used as a marker for the recycling endosome,^59–61^ we found no statistically significant difference in colocalization with transferrin (Fig. 4c).

As expected, we find intracellular insulin trafficking to be cell type dependent,^2,11,65^ with different diffusion and colocalization patterns found in cancer cells (HeLa) as compared to HEK293 (Fig 2e, Fig 4c and supplementary Fig. 13). In HeLa cells, we do not report a significant difference in intracellular diffusion with DeepSPT nor a difference in endolysosomal colocalization with Colocalization Fingerprinting (see Supplementary Fig. 13). These results are consistent with insulin signaling and degradation known to be highly tissue and receptor isoform-specific, and reliant on variation in receptor isoform populations.^11^

## Discussion

IAsp, is one of the insulin analogs that structurally resembles HI the closest, however it does not fully mimic the uptake, secretion, and cellular response of endogenous insulin.^56,66^ IAsp’s point mutation from proline to aspartic acid at position B28, is placed outside the primary receptor binding region, to maintain structural similarity to HI while destabilizing the dimer and hexamer oligomers, thus promoting rapid cellular uptake of monomers from the bloodstream.^67,68^ Although, HI and IAsp display similar receptor affinities to both the insulin and the IGF I receptors at physiological pH, slight differences in receptor off-rates^10,64^ and insulin receptor isoform activation have been reported, consequently impacting downstream Protein Kinase B (AKT) and Extracellular Signal-Regulated Kinase (ERK) activation.^11^ While alterations in receptor binding are expected to lead to different cellular responses, which may have important implications, especially for prolonged treatments,^69,70^ the cellular and molecular mechanisms causing variations in uptake, secretion and response are not fully understood and the full signaling range and trafficking characteristics of insulin are still being uncovered. We firmly stress that IAsp is safe in clinical use and, along other fast-acting analogs, vital for patients who need rapid insulin response postprandial, imperative for patients suffering from the type 1 diabetes.^66,71^

Studies on internalization, signaling and trafficking pathways of insulin are rare and primarily based on bulk receptor binding assays conducted in non-native conditions such as purified receptors, serum-starved, dormant or dead cells, cold conditions or fixed pH.^6–10^ These conditions can perturb or disrupt endocytosis and possibly impact receptor binding properties, which can change both insulin and analogs internalization as well as their cellular response ^9,22,72^

To study the cellular trafficking pathways of HI^655^ and IAsp^655^ in physiologically relevant conditions, we used live-cell imaging and single-particle tracking at native insulin receptor levels. Precise quantification of differences between the two isoforms in their intracellular motion and colocalization propensities in the endolysosomal pathway requires spatiotemporally resolving endolysosomal and recycling compartments. However, the crowded cellular milieu, highly heterogeneous diffusion, and experimental noise present a formidable challenge for current tools to estimate colocalization with intracellular compartments, obscuring the quantification of intracellular trafficking and sorting mechanisms.^31–33,37,38,48,49,54,55^

To overcome this, we introduced a combined machine learning framework, DeepSPT and Colocalization Fingerprinting that utilize diffusional analysis to decipher colocalization in crowded environments with enhanced precision. Utilizing spatiotemporal behavior improves the reliability of colocalization analysis and expands colocalization output from a single value to a time-resolved distribution, maintaining information on particle heterogeneity and temporal variations. The reliable quantification of this method revealed a significant difference in intracellular diffusion rates between insulin variants, with IAsp^655^ displaying a 50% larger median average step length compared to HI^655^. This is consistent with colocalization fingerprints revealing increased colocalization with the slow-moving lysosomal compartments for HI^655^ as compared to IAsp^655^. Together, they underline the strength of Colocalization Fingerprinting, which convincingly separates and quantifies endosomal localizations of insulin analogs varying by a single point mutation, shown to profoundly alter intracellular trafficking and colocalization propensities.

We selected kidney cells (HEK293) due to their role in insulin clearance in diabetic individuals, as subcutaneous injection circumvents initial hepatic clearance.^65,73^ The differences in intracellular trafficking of IAsp and endogenous insulin in HEK293 could thus link to largely unresolved cellular clearance and degradation pathways of both recombinant insulin and analogs. The direct recording here suggests that the non-endogenous pharmacokinetic and pharmacodynamic properties may be partially due to differences in intracellular trafficking. Moreover, we find intracellular trafficking to be cell type dependent, with different diffusion and colocalization patterns found in cancer cells (HeLa) as compared to HEK293 (see Supplementary Fig. 13, Fig. 2e and 4c). These trafficking differences likely stem from cell type and receptor isoform-specific insulin uptake, as kidney cells mainly facilitating insulin clearance whereas cancer cells rely on insulin for glucose uptake and enhanced cell growth.^2,11,65^

The high fidelity SPT combined with the DeepSPT and Colocalization Fingerprinting toolbox tackles the challenges of analyzing intracellular colocalization in crowded environments. Leveraging this framework on insulin we demonstrate its sensitivity to robustly detect and quantify even minute hitherto unknown differences in intracellular trafficking for insulin analogs varying by a single point mutation, and detect cell type dependent differences.While the full implications of these differences in intracellular insulin behavior are subject of current studies, the data are consistent with our recently introduced notion that spatiotemporal motion links to biological function. This could have implications for drug efficacy and thus inform pharmacodynamics and pharmacokinetics, guiding the development of new future therapeutics. The reliable analysis toolbox for intracellular trafficking and sorting, would be key to enhance our understanding on how modifications at the molecular level impact cellular response, for a wide range of biological systems. Thus, the method is of great interest for various biological applications where cellular internalization, intracellular trafficking and endosomal localizations are important, from endosome trafficking to viral entry, drug delivery and endosomal escape.

## Methods

### Experimental

#### Materials

All chemicals are of analytical grade and purchased from Sigma-Aldrich Denmark, unless otherwise stated. Atto655-NHS ester was purchased from ATTO-TEC. All cellular compartment labels; CellLight^TM^ Early Endosome-GFP, BacMam 2.0, CellLight^TM^ Late Endosome-GFP, BacMam 2.0, LysoTracker® Green DND-26 and Transferrin-AlexaFluor^TM^488 were purchased from Invitrogen^TM^. Recombinant human insulin was purchased from Thermo Fisher USA. Insulin aspart (NovoRapid®, Novo Nordisk) was of medical grade and purchased from a local pharmacy. MilliQ water was used for all aqueous preparations. 10mM HBSS-HEPES buffer pH 7.4 without Ca^2+^ and Mg^2+^ was used as an imaging buffer for all experiments.

#### Insulin preparations

Human Insulin (HI) and Insulin aspart (IAsp) were conjugated to Atto655 at the lysine at position B29 by the same protocol, following previous publications.^74,75^. Prior to conjugation, IAsp was purified using a Biotage Isolera instrument. Human insulin (21 mg, 3.6 μmol, 3 equivalents) or freshly purified insulin aspart (15 mg, 2.6 μmol, 2 equivalents) was suspended in 0.1 M Tris Buffer (0.2 mL), and pH was adjusted to 10.5 to dissolve it completely. Atto655-NHS ester (1.0 mg, 0.00122 mmol, 1.0 equivalent for HI) (1.1 mg, 0.00182 mmol, 0.70 equivalents for IAsp) was dissolved in DMF (0.3 mL), added dropwise over 5 minutes (HI) or 3 minutes (IAsp) to the stirred solution of Insulin, and the reaction mixture was allowed to stir for 15 min. The reaction was monitored by LCMS.^76,77^ The reaction mixture was subsequently diluted with 2.0 mL (HI) or 1.5 mL (IAsp) H_2_O and pH was adjusted to pH 7.8. The products were isolated using RP-HPFC (Biotage Selekt), applying Biotage SNAP ultra-column (C18, 30 g, 25 µm). CH_3_CN/H_2_O mixed with 0.1% formic acid was used as eluent at a linear gradient of 5-50% CH_3_CN over 20 min and a flow rate of 25 mL/min. Each fraction was analyzed through LCMS (for LysB29Atto655-HI we refer to previous published results^74^, for LysB29Atto655-IAsp see Supplementary Fig. 1 and Table 1). Monosubstituted products were collected separately, CH_3_CN was removed at reduced pressure using a rotary evaporator, followed by lyophilization to give a dark green (or blackish green) powder as a product (LysB29Atto655-HI Yield: 5.6 mg, 79%) (LysB29Atto655-IAsp Yield: 4.2 mg, 51%). High-resolution mass spectrometry (HR-MS) was performed on an ultra-high pressure liquid chromatography Dionex Ultimate 3000 instrument (Thermo Scientific) coupled to a Bruker Impact HD II quadrupole time-of-flight mass spectrometer (QTOF). Purification of conjugates was performed on the Biotage-Isolera HPFC 300 system with a C18 column (SNAP Ultra, C18, 30 g).

#### Cell culture

HEK293 cells were obtained from Cell Line Service (CLS, now Cytion, received at split level 27) and cultured in Dulbecco’s Modified Eagle Medium (DMEM) with 10% Fetal Bovine Serum (FBS) at 37°C with 5% CO_2_. Experiments were performed on cells with split levels between 29 and 37. HeLa cells were kept under the same culturing conditions.

#### Sample preparation

Prior to imaging, 20.000 HEK293 cells were seeded in Ibidi IbiTreat 8-well plates and cultured at 37°C and 5% CO_2_ for 2 days. For early and late endosome co-labeling, 1 μL/10.000 cells (3μL per well) CellLight^TM^ Early Endosome-GFP BacMam 2.0 or CellLight^TM^ Late Endosome-GFP BacMam 2.0 was added respectively during incubation, 16-22 hours prior to imaging. Lysosomes were labeled by LysoTracker® Green DND-26 in a 1:20.000 dilution in the imaging buffer and added during incubation, 1 hour before imaging. The recycling pathway was probed by Transferrin-Alexa Fluor^TM^ 488 conjugate by incubating 5μg per well for 1 hour prior to imaging. Insulin, either 0.05 mg/mL HI-Atto655 or 0.05 mg/mL IAsp-Atto655, was added to the HEK293 cells 1 hour prior to imaging in the imaging buffer. For each condition, one insulin analog was added and one cellular compartment tagged to follow their diffusion in real time using Spinning Disc Confocal Microscopy (SDCM), yielding a total of eight conditions (2 insulin variants and 4 cellular compartments, see Figure 1a). The same overall protocol was followed for HeLa cells, with 10.000 cells seeded per well.

### Live cell SDCM microscopy

#### Microscopy

All experiments were conducted on an inverted Spinning Disk Confocal Microscope (SDCM) (Olympus SpinSR10) equipped with an oil immersion 60X objective (Olympus) with a numerical aperture of 1.4 and a CMOS camera (photometrics PRIME 95B) with an effective pixel size of 183 nm x 183 nm. Imaging was performed with a dual-imaging module (Hamamatsu Photonics W-VIEW GEMINI A12801-01). Insulin analogs with Atto655 fluorophores were excited using a 640 nm laser line and imaging was performed with 100% laser power. Compartment labels; Rab5a-EmGFP (CellLight^TM^ Early Endosome-GFP, BacMam 2.0), Rab7a-EmGFP (CellLight^TM^ Late Endosome-GFP, BacMam 2.0), LysoTracker (LysoTracker® Green DND-26) and Transferrin-AlexaFluor^TM^488 were excited using a 488 nm laser line, with 10%, 10%, 3% and 10% laser power respectively. Lasers were aligned manually prior to all experiments with dual-labeled liposomes and the field of view was adjusted to minimize chromatic aberration. All videos were recorded with 3 EM gain and 30,4 ms exposure time for a total frame rate of 36 ms including lag-time, in SDCM streaming settings for 2000 frames at 37°C. HEK293 cells were washed once with fresh, pre-heated 10mM HBSS-HEPES buffer pH 7.4 without Ca^2+^ and Mg^2+^ immediately before imaging each technical replicate. HeLa cells followed the same overall protocol but used laser powers 20%, 20%, and 3% for early endosomes, late endosomes, and lysosomes, respectively (transferrin not measured). Each condition was performed in 2 biological replicates and imaged in 3 technical replicates with 2 videos taken for each well, yielding a total of 12 videos per condition.

### Image analysis

#### Tracking

Single particle tracking of vesicles containing insulin analogs or intracellular compartments was performed with a previously described in-house tracking script based on TrackPy.^78–81^ The particles were identified using the following tracking parameters; object diameter of 9 pixels, search range of 5 pixels, gap closing (memory) 1 frame, and with mean-multiplier between 0.6 and 0.9 evaluated manually for each biological replicate. Only trajectories longer than 20 frames were used for consecutive analysis.

#### Background correction

Post-processing of all compartments (early endosomes, late endosomes, and lysosomes) and transferrin trajectories were completed with hard thresholds, keeping particles with mean eccentricity < 0.3 and mean corrected integrated intensity > 0. For HI-Atto655, the thresholds mean eccentricity < 0.2 and mean corrected integrated intensity > 0 were used. Background correction for IAsp-Atto655 trajectories was performed by training a logistic regression model with Scikit-Learn in videos with or without the insulin analog, using 8 detection features (mean step length, track duration, diffusion coefficient, eccentricity, size, mass, ellipticity, and signal) to filter detections.^82^

#### Track densities

To estimate experimentally observed densities (tracks/μm^2^) of endolysosomal and recycling compartments, the number of trajectories localized within five 10×10 μm boxes were counted per condition, revealing 3-20 tracks per area resulting in densities of 0.03-0.15 tracks/μm2. Most often ∼10 tracks were observed per 10×10 μm area, also seen with the mean ± sem of 10.13 ± 0.64 detections.

### DeepSPT and Colocalization Fingerprinting Analysis

#### DeepSPT

The intracellular diffusion of biomolecules is highly heterogeneous with large spatiotemporal variation. To analyze the diffusional properties and temporal behavior of insulin and cellular compartments, a deep learning-assisted framework for single particle tracking analysis, termed DeepSPT (Kæstel-Hansen et al.^58^), was utilized. DeepSPT is designed for extracting information from heterogeneous single-particle diffusion and consists of three modules: 1) temporal segmentation of diffusional behavior, 2) diffusional fingerprinting, and 3) a task-specific classifier.

#### Diffusional featurization by DeepSPT

Utilizing the first two modules of DeepSPT^58^, all trajectories were temporally segmented to extract temporal diffusional behavior and subsequently transformed into comprehensive feature representations consisting of over 40 diffusional features. Specifically, the temporal segmentation module semantically segments each time point of each trajectory into one of four diffusional behaviors (Brownian, directed, confined, or subdiffusive motion). Subsequently, the diffusional fingerprinting module, utilizing these semantic predictions of temporal behavior and the diffusional of each trajectory, constructs a unique fingerprint per trajectory, based on particle motion.

#### Classifying endosomal diffusion using DeepSPT

Leveraging the entire DeepSPT pipeline, we trained an application-specific classifier to differentiate endosomal identity solely utilizing diffusional characteristics. Specifically, we trained a random forest classifier with a max depth of 10 and otherwise default scikit-learn parameters on diffusion observed for different endosomal compartments.^82^ This classifier was trained on endosomal tracks longer than 50 time points yielding 18980, 17725, and 13859 single-particle trajectories of early endosome, late endosome, and lysosome, respectively. The classifier was evaluated in a ten-fold cross-validation scheme both in the three-class problem of differentiating the three compartments and in the two-class problem of differentiating early endosomes and late endosomes from lysosomes. We tasked the classifier with categorizing trajectories of insulin variants with durations over 50 time points to the compartment with the most similar diffusion, thus investigating diffusional similarity for 11925 and 17457 trajectories for HI^655^ and IAsp^655^, respectively.

#### Simulated colocalization

To train and validate the colocalization model, actual colocalized trajectories and false colocalized trajectories with close proximity was simulated using the diffusion simulation framework underlying the method in Kæstel-Hansen et al.^58^, to construct heterogeneously diffusing trajectories with widely distributed diffusional characteristics. For true colocalization, simulated trajectories are duplicated and these duplicated pairs represent actual colocalization. To account for localization errors, technical artifacts, and to mask similarities, the tracks are translated randomly in x and y, and perturbed by Gaussian noise to each position, in each trajectory, independently. For false colocalization, two simulated trajectories were randomly drawn and trimmed to match trajectory durations to the shortest. Displacements of the trajectories are scaled to ensure comparable diffusion coefficients to add further complexity. In addition, the two trajectories are perturbed by Gaussian noise, and only time points within a user-defined distance are kept, subsequently, the two trajectories are randomly translated in x and y. The result is a pair of spatially proximate trajectories with uncoupled diffusion to represent false colocalization. To increase trajectory similarity for false colocalization pairs, a subset of pairs is drawn, based on a user-defined probability, and aligned by Procrustes superimposition. The distribution of simulated trajectory lengths is included in Supplementary Fig. 2 and examples of simulated trajectories are in Supplementary Fig. 3.

#### Pipeline for the initial colocalization module (M1)

Segments of colocalization are initially defined based on temporally consistent spatial proximity between recorded trajectories across imaging channels. The lock-step Euclidean distances are computed between all trajectories in the primary channel and in the secondary channel. If a user-defined number of consecutive frames (“min_coloc”) is below a user-defined search distance (“distthreshold”) the track segments are defined as colocalizing. Noise and localization errors can produce sudden spikes in inter-track distances, which can break a true colocalization segment. To mitigate this, a user-defined number of frames is permitted outside the defined search distance (“forgiveness”), and colocalizing segments on either side of this spurious loss of colocalization will be connected. To increase certainty in identified segments being true colocalization and minimizing registered spurious proximity multiple filters are implemented: minimum required total colocalization length, minimum required average lock-step Euclidean distance, and a minimum required Pearson correlation between identified segments. Importantly, the identified segment pairs of colocalization are subjected to colocalization fingerprinting (M1 and M2) and evaluated for true colocalization by the trained colocalization fingerprint classifier.

#### Colocalization fingerprint (M2)

Colocalization between two biological entities is typically identified by spatial proximity or by temporal consistency of proximity. To enhance the differentiation of true colocalization versus spurious proximity we establish colocalization fingerprinting to combine spatial proximity with similarity of diffusion between the two entities. For a pair of trajectories suspected to colocalize, both trajectories are transformed into a set of 23 selected features.

Firstly, DeepSPT is utilized to establish diffusional feature characteristics for the pair of trajectories. 15 out of ∼40 diffusional features were selected. These include anomalous diffusion exponent, diffusion coefficient from time-averaged mean square displacement (MSD), quality of the fit to the MSD curve, efficiency, log efficiency, fractal dimension, MSD ratio, average step length, average MSD, maximum step length, difference in minimum and maximum step length, area of trajectory, number of changepoints in diffusional behavior, and the apparent diffusion coefficient.^58,83^ The selection of features was based on maximizing downstream prediction accuracy and visual inspection of features for manually identified colocalizing trajectories in experimental data. One of these 15 features is made by combining 4 original DeepSPT features to the respective maximum value to minimize zero value features, each of the four original features describes the time a trajectory expresses a given diffusional behavior (Brownian, directed, confined, or subdiffusive).^58^

Secondly, eight additional trajectory similarity features were implemented into the framework. 1: Partial Curve Mapping between the two trajectories. 2: Bray-Curtis dissimilarity between the two trajectories. 3: Cross-correlation between the two trajectories in x and y, independently. 4: Normalized dot-product of the vectors formed between each track’s start and end position. 5: Cosine similarity for the vectors formed by each trajectory’s start and end position. 6: Divergence measure, defined as the distance between start points of the trajectories versus endpoints relative to the standard deviation of all distances. Gaussianity of inter-trajectory distances is defined as the 95% quantile distance relative to the expected 95% confidence interval for Gaussian spread (1.96 times the standard deviation of distances). Combined these 23 features form the colocalization fingerprint of two trajectories.

#### Colocalization fingerprinting classifier (M3)

To evaluate colocalization using the colocalization fingerprint, a supervised machine-learning model is trained on simulated colocalization, collected using the described simulation protocol. A random forest classifier is utilized with a max depth of 2 and otherwise default scikit-learn parameters.^82^ During training simulated true colocalizing track durations are resampled to match the length distribution and sample size of false colocalization to mitigate model bias (see Supplementary Fig. 1). Classification performance is evaluated using a ten-fold, shuffled cross-validation scheme.

#### Stress test of colocalizing segments’ duration

Trajectories are simulated following the described simulation protocol with the additional requirement of fixed trajectory durations of 5, 7, 10, 20, 50, 100, or 200 time points. Each condition includes 1000 trajectories and for each condition classification performance is evaluated using a ten-fold, shuffled cross-validation scheme.

#### Stress test of colocalizing segments’ similarity to concerted motion

Trajectories are simulated following the described simulation protocol for true colocalization, however, for false colocalization only pairs with a specified cosine similarity to concerted motion are kept. This is implemented by measuring the cosine similarity between the 23-dimensional colocalization fingerprint representation of a pair and the colocalization fingerprint for perfectly concerted motion. Here, similarities of 0.99, 0.9, 0.7, and 0.6 with a tolerance of 0.02 are accepted. The simulation protocol runs until a minimum of 1000 false colocalization pairs with the specified similarity are identified. To mitigate class imbalance, true colocalization pairs are randomly sampled with a sample size identical to the false colocalization pairs. Classification performance is evaluated using a ten-fold, shuffled cross-validation scheme.

#### Stress test of increasing population densities

Heterogeneously diffusing trajectories that randomly sample Brownian, directed, confined, and subdiffusive motion are simulated using the framework from Kæstel-Hansen et al.^58^ Diffusional characteristics are widely distributed and convolved with localization errors as per Kæstel-Hansen et al. with a few exceptions.^58^ Specifically, track duration is fixed at 500 time points, the diffusion coefficient is lognormally sampled from 0.1 to 0.5 μm^2^/s, and trajectory positions are sampled corresponding to a 30 ms frame rate. To compare, SPT experiments used 36 ms frame rates and observed 0.13 and 0.18 μm median step lengths, yielding apparent diffusion coefficients of 0.12 to 0.225 μm^2^/s (using the MSD, 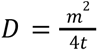, where *m* is the median step length, t is the frame rate, and D is the diffusion coefficient), respectively. These trajectories are initiated at random coordinates within a 20×20 μm box with reflective boundaries, meaning trajectories are reflected when reaching the box limits (see Supplementary Fig. 4). Subsequently, the first module for initial colocalization by Colocalization Fingerprinting is run with parameters: minimum number of consecutive frames of 5, a search distance threshold of 0.5 μm, and forgiveness of 3 frames. Pearson correlation threshold of 0.1 (Supplementary Fig. 5) or 0.5 (Fig. 3d), minimum total colocalizing length of 5 frames, and a minimum average Lock-step Euclidean distance of 0.5 μm. M1 is executed on the simulated trajectories to propose colocalizing segments. Every proposed segment corresponds to an identification by solely using a distance-based approach. For implementing more stringent approaches, each proposed colocalizing pair is evaluated by either requiring Pearson’s correlation above 0.5 or by executing the subsequent modules of Colocalization Fingerprinting (M2 and M3).

#### Colocalization settings for dual-color SPT

The settings used for initial colocalization identification included a minimum insulin trajectory length of 50 time points, a minimum number of consecutive frames of 3, a search distance threshold of 3 μm, and forgiveness of 7 frames. Pearson correlation threshold of 0.1, minimum total colocalizing length of 20 frames, and a minimum average Lock-step Euclidean distance of 1.85 μm. Settings were purposefully sat leniently to account for noise and any interchannel aberrations. In the rare case of an insulin trajectory having multiple proposed partners, the proposal with the longest colocalization duration weighted by a probability of colocalization predicted by Colocalization Fingerprinting is selected.

## Data and code availability

Data and code will be made fully available upon publication on ERDA.ku.dk and Github.

## Author contributions

S.V.B., J.K.H., and N.S.H. wrote the manuscript with input from all authors. S.V.B. and A.J.N. performed experimental design, sample preparation, and cell culturing. S.V.B. carried out fluorescence imaging and SPT in HEK293. S.V.B. and A.J.N. executed imaging and SPT in HeLa. S.V.B. conducted fluorescence image and SPT analysis. K.J.J. and N.K.M. synthesized and formulated the labeled insulin compounds. J.K.H. developed and evaluated Colocalization Fingerprinting, the simulation framework, and employed DeepSPT. S.V.B and J.K.H. conducted statistical and diffusional analysis. S.V.B., J.K.H., and N.S.H. performed data interpretation. N.S.H. designed the project and performed overall project management.

## Acknowledgements

We thank the members of our laboratories for helpful discussions and encouragement. We thank Shunliang Wu for his contributions to insulin preparation and characterization. This work was funded by the Villum Foundation (grant 18333 and 40801), The Novo Nordisk foundation challenge center for Optimised Oligo Escape and Control of Disease (NNF23OC0081287), and The Center for 4D Cellular Dynamics (NNF22OC0075851). N.S.H. is affiliated with the Novo Nordisk Foundation Center for Protein Research (CPR) funded by a generous donation from the Novo Nordisk Foundation (grant NNF14CC0001). S.V.B, J.K.H., A.J.N., K.J.J. and N.S.H are members of the Integrative Structural Biology Cluster (ISBUC) at the University of Copenhagen.

## List of Abbreviations

HI: Human Insulin
HI^655^: Atto655 labeled Human Insulin
IAsp: Insulin Aspart
IAsp^655^: Atto655 labeled Insulin Aspart
HEK293: Human embryo kidney 293 cells
DeepSPT: Deep learning-assisted single-particle diffusional analysis
SPT: Single-particle tracking
EE: Early endosomes
LE: Late endosomes
LS: Lysosomes
TF: Transferrin
IGF-1: Insulin-like growth factor 1
AKT: Protein Kinase B
ERK: Extracellular signal regulated kinase
MSD: Mean square displacement

