## Supplementary information for "Diverse Intracellular Trafficking of Insulin Analogs by Machine Learning-based Colocalization and Diffusion Analysis"

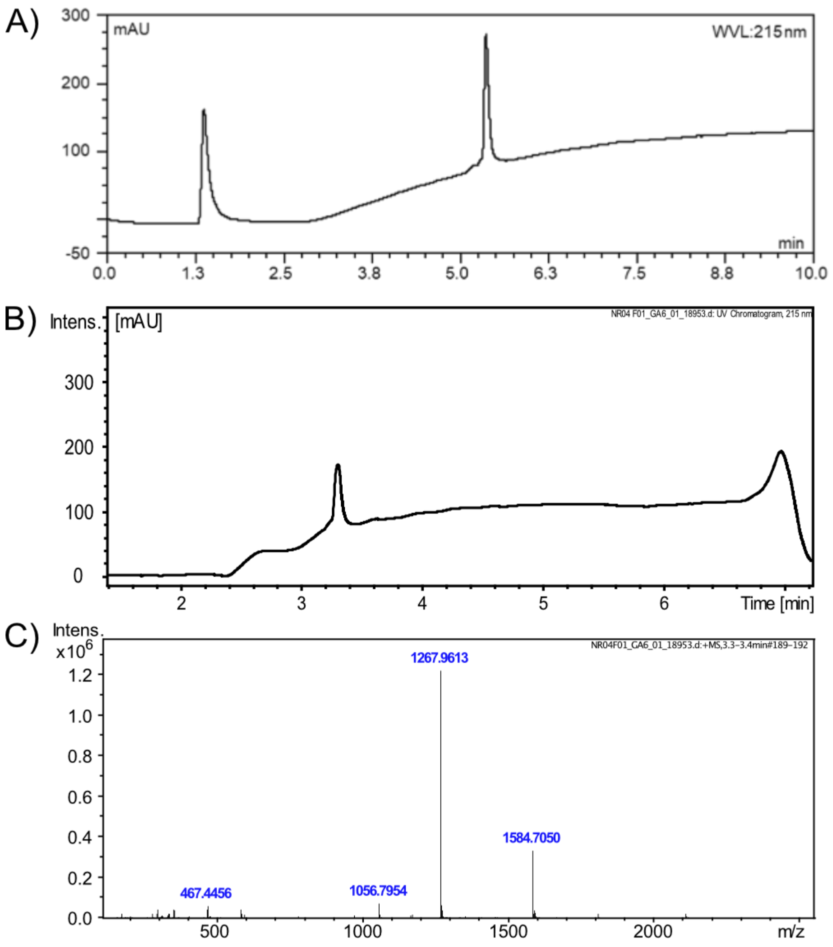


**Supplementary Figure 1: A)** Low resolution LCMS chromatogram of purified Atto655 labeled Insulin aspart (LysB29-Atto655-Insulin aspart). The high resolution LCMS chromatogram and mass spectra are shown in **B)** and **C)** respectively.

**Supplementary Table 1:** High resolution LCMS analysis of LysB29-Atto655-Insulin aspart. Molecular formula: $C_{256}H_{381}N_{65}O_{79}S_6$

| Serial No. | Peak | Calculated | Observed |
|---|---|---|---|
| 1. | $[M]^+$ | 6331.82 | Not Observed |
| 2. | $[M+4H]^{4+}$ | 1584.46 | 1584.71 |
| 3. | $[M+5H]^{5+}$ | 1267.77 | 1267.96 |
| 4. | $[M+6H]^{6+}$ | 1056.64 | 1056.80 |

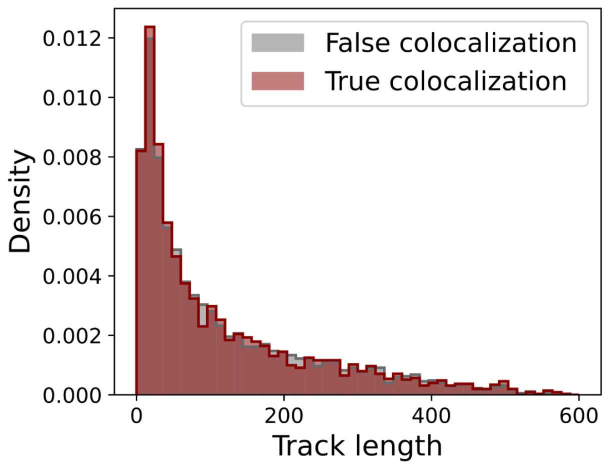


**Supplementary Figure 2:** Simulated trajectory durations for true and false colocalization. Distributions are balanced to mitigate model bias.

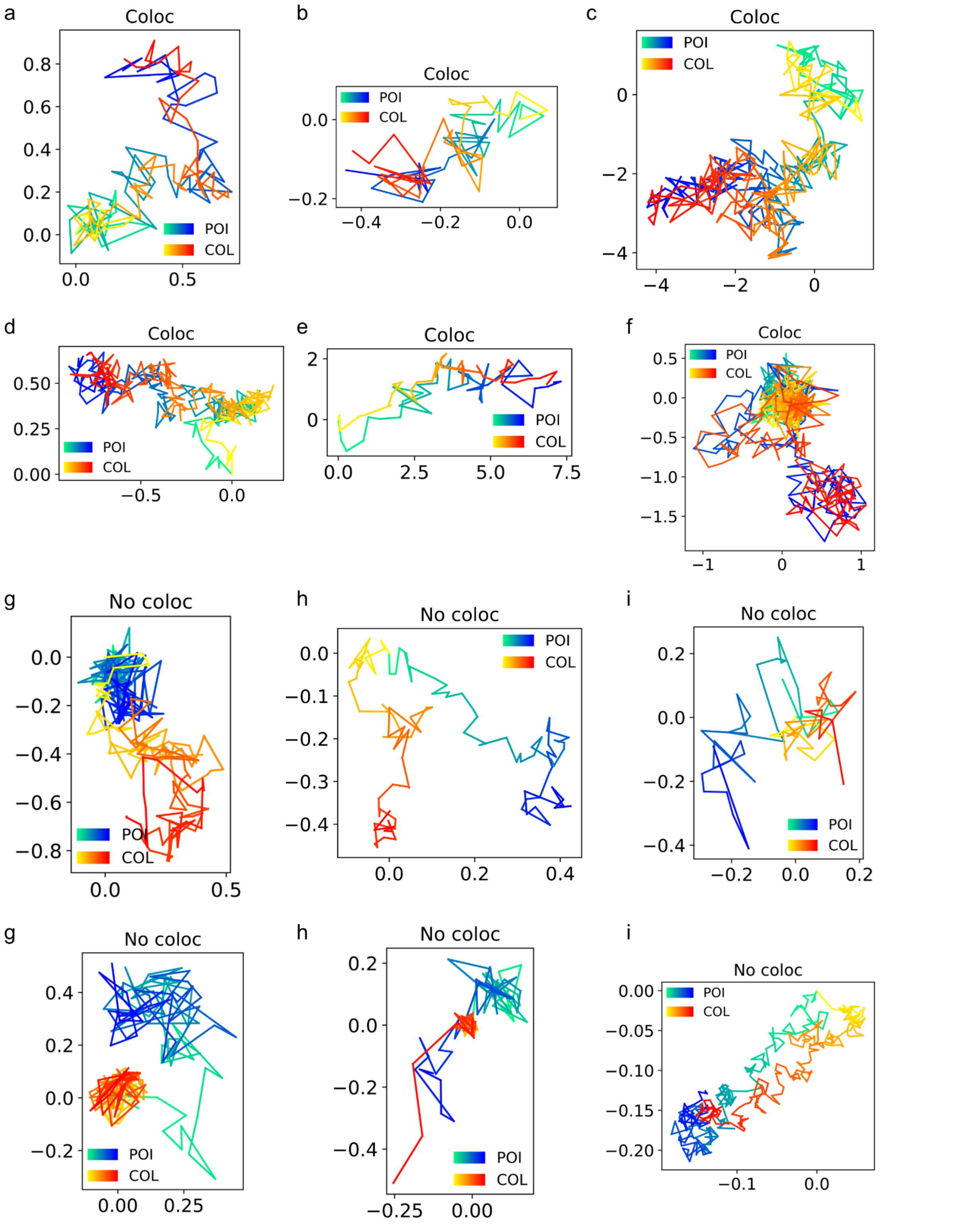


**Supplementary Figure 3:** Examples of simulated trajectories for true (Coloc) and false (No coloc.) colocalization. Trajectories are colored reflecting the temporal axis from light to darker shades. All coordinates are in micrometers. POI: particle of interest, i.e., the track that had its colocalization evaluated. COL: identified colocalization partner.

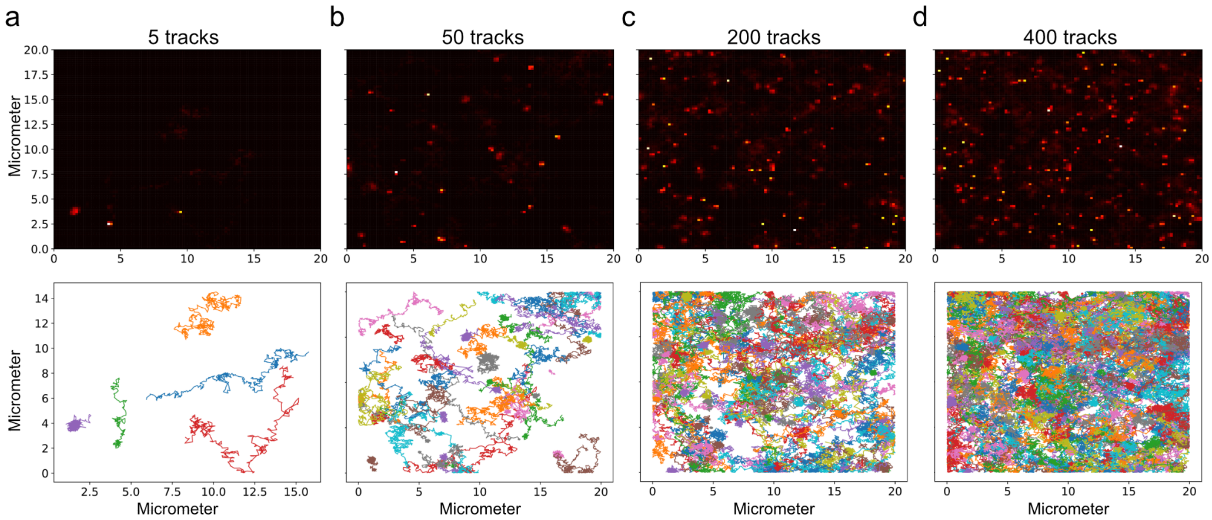


**Supplementary Figure 4:** Simulated data of an enclosed space (20x20 µm) with reflective boundary conditions and an increasing number of tracks to mimic increasingly crowded environments. Top row: 2D histograms displaying spatial density of localizations. Bottom row: Showing trajectories. **a)** 5 tracks (0.0125 tracks/µm$^2$). **b)** 50 tracks (0.125 tracks/µm$^2$). **a)** 200 tracks (0.5 tracks/µm$^2$). **b)** 400 tracks (1 track/µm$^2$).

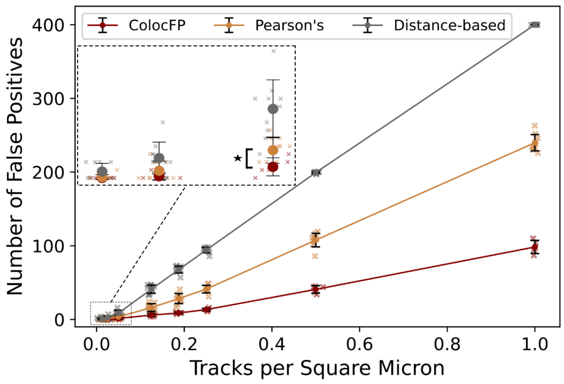


**Supplementary Figure 5:** Evaluation of Colocalizational fingerprinting (with the more lenient filter requiring Pearson's correlation above 0.1) and benchmark approaches in filtering out spurious colocalization in an enclosed space (20x20 µm) with an increasing density of non-interacting particles (see Supplementary Fig. 3), hence no true colocalization. The number of identified colocalizing segments directly denotes the false positive rate and the susceptibility to crowding. Zoom-in depicts the low-density regime for visual clarification. Colocalizational fingerprinting consistently identifies significantly fewer false positives compared to Pearson's down to densities of 0.05 tracks/µm$^2$ (20 tracks per 400 µm$^2$), as evaluated by a two-sided Welch's t-test (p-value<0.001, N=10 per approach). Errors depict standard deviation across ten independent crowding simulations (see Methods).

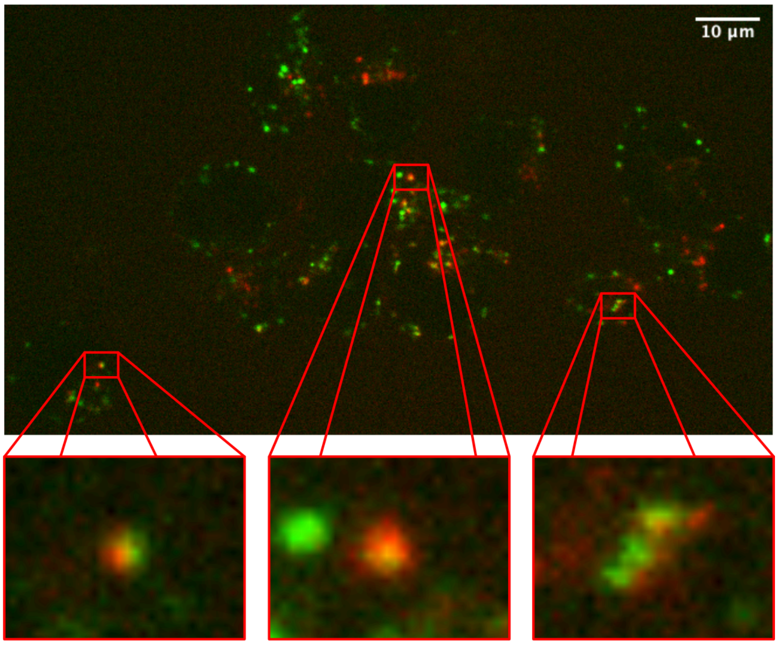


**Supplementary Figure 6:** Chromatic aberrations in spinning disk confocal microscopy produce wavelength-dependent distortions in position.[1–3] Top: Cropped field of view. Bottom left: Zoom-in showing a colocalized signal from the left side of the field of view, displaying a shift in the red light to the left. Bottom middle: Zoom-in from the center of the field of view showing a green signal and a colocalized (red and green) signal in the center of the zoom-in with minimal spatial distortion. Bottom right: Zoom-ins, from the right side of the field of view displaying red colors shifted to the right. Notably, the chromatic aberration increases radially and differs depending on the position relative to the center of the field of view. [1,2] The chromatic offset reported here reaches up to ~6 pixels by careful visual inspection, i.e., ~1200 nm. Importantly, the field of view is cropped around the center to avoid even larger chromatic offsets further from the center. Human Insulin (red), lysosomes (green). Scale bar 10 µm.

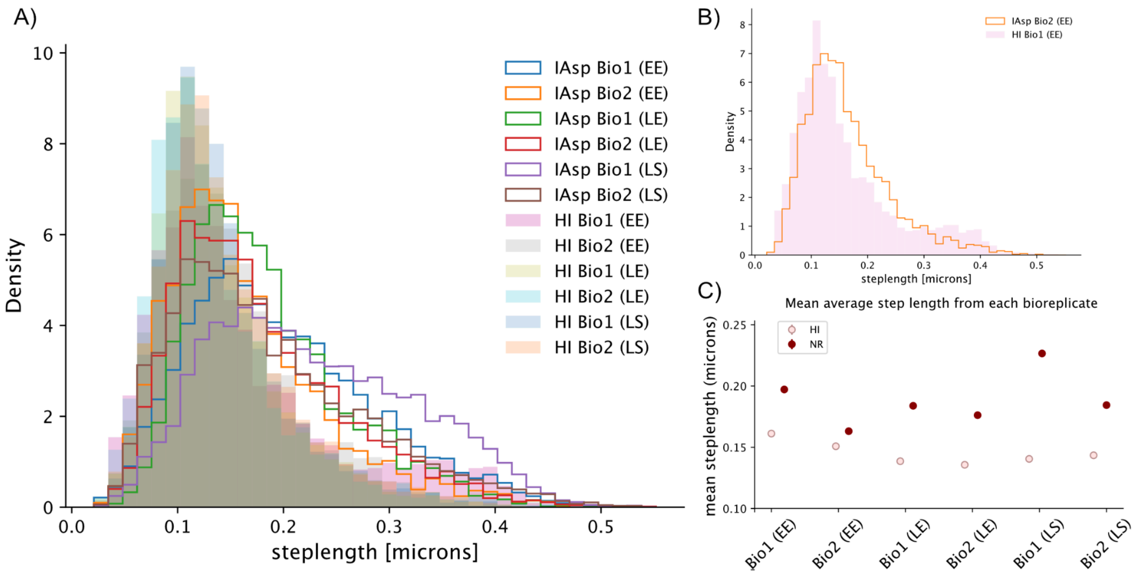


**Supplementary Figure 7:** Insulin mean step length analysis. **a,** Average step length distributions of trajectories for each bio replicate from each experimental condition of human insulin (HI, full) and insulin aspart (IAsp, lines). A one-sided Welch's t-test found IAsp to have significantly longer average mean step lengths compared to HI for all bio replicates except one (See Supplementary Table 1). **b,** Shows the comparison between the two samples (IAsp orange line, HI pink) which did not show a significant difference. The distribution of IAsp is seen to generally be moved towards higher mean step lengths with HI showing a secondary population with higher mean step lengths, not present in the remaining HI samples. **c,** Mean average step lengths of each bio replicate. In all cases, IAsp shows higher average step lengths. Errors depict the standard error on the mean. Number of particles in each bio replicate: $N_{HI\ EE\ Bio1} = 2280$ , $N_{HI\ EE\ Bio2} = 3342$, $N_{HI\ LE\ Bio1} = 3487$, $N_{HI\ LE\ Bio2} = 2089$, $N_{HI\ LS\ Bio1} = 5841$, $N_{HI\ LS\ Bio2} = 3662$, $N_{IAsp\ EE\ Bio1} = 7269$, $N_{IAsp\ EE\ Bio2} = 3024$, $N_{IAsp\ LE\ Bio1} = 4692$, $N_{IAsp\ LE\ Bio2} = 4756$, $N_{IAsp\ LS\ Bio1} = 10976$, $N_{IAsp\ LS\ Bio2} = 4885$. EE = Early endosomes, LE = Late endosomes, LS = lysosomes.

**Supplementary Table 2:** Calculated p-values for the difference in mean average step lengths between Human Insulin (HI) and Insulin aspart (IAsp) from one-sided Welch's t-test with the null hypothesis the mean average step length of HI is greater than IAsp, compared between all bio replicates. All comparisons but one show significant differences. For raw mean step length distributions for each bio replicate, see Supplementary Fig. 6. Number of samples in each replicate: $N_{HI\ EE\ Bio1} = 2280$, $N_{HI\ EE\ Bio2} = 3342$, $N_{HI\ LE\ Bio1} = 3487$, $N_{HI\ LE\ Bio2} = 2089$, $N_{HI\ LS\ Bio1} = 5841$, $N_{HI\ LS\ Bio2} = 3662$, $N_{IAsp\ EE\ Bio1} = 7269$, $N_{IAsp\ EE\ Bio2} = 3024$, $N_{IAsp\ LE\ Bio1} = 4692$, $N_{IAsp\ LE\ Bio2} = 4756$, $N_{IAsp\ LS\ Bio1} = 10976$, $N_{IAsp\ LS\ Bio2} = 4885$. ** p-value < 0.01

| HI / IAsp | EE Bio1 | EE Bio2 | LE Bio1 | LE Bio2 | LS Bio1 | LS Bio2 |
|---|---|---|---|---|---|---|
| EE Bio 1 | ** | ** | ** | ** | ** | ** |
| EE Bio2 | 0.20 | ** | ** | ** | ** | ** |
| LE Bio1 | ** | ** | ** | ** | ** | ** |
| LE Bio2 | ** | ** | ** | ** | ** | ** |
| LS Bio1 | ** | ** | ** | ** | ** | ** |
| LS Bio2 | ** | ** | ** | ** | ** | ** |

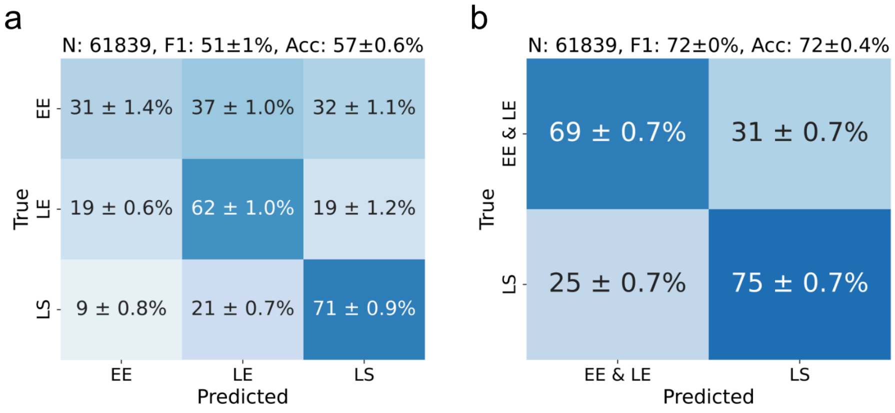


**Supplementary Figure 8:** Confusion matrix of classifying endosomal compartments from diffusion alone using DeepSPT (see Methods).[4] **a,** Three-class prediction task of early endosome (EE), late endosome (LE), and lysosome (LS). Additional classification metrics accuracy (Acc.), F1-score (F1), number of tracks (N). **b,** Two-class prediction task by collapsing early and late endosomes under the same label (EE & LE). All errors depict standard deviation across a ten-fold cross-validation scheme.

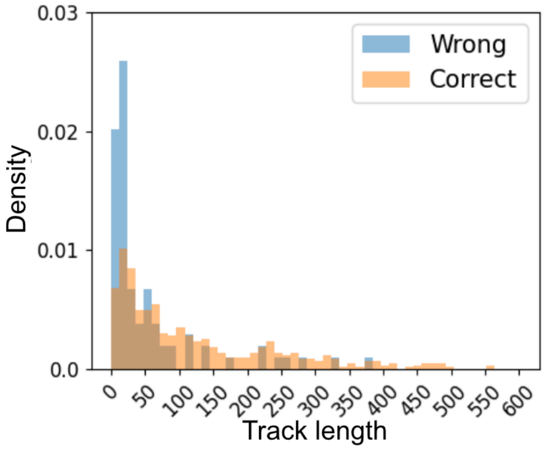


**Supplementary Figure 9:** Distributions of trajectory length for simulated trajectories correctly and incorrectly predicted by Colocalizational Fingerprinting. Incorrectly predicted trajectories are predominantly short, while correctly predicted tracks span all track lengths. This is expected as short trajectories contain less information.

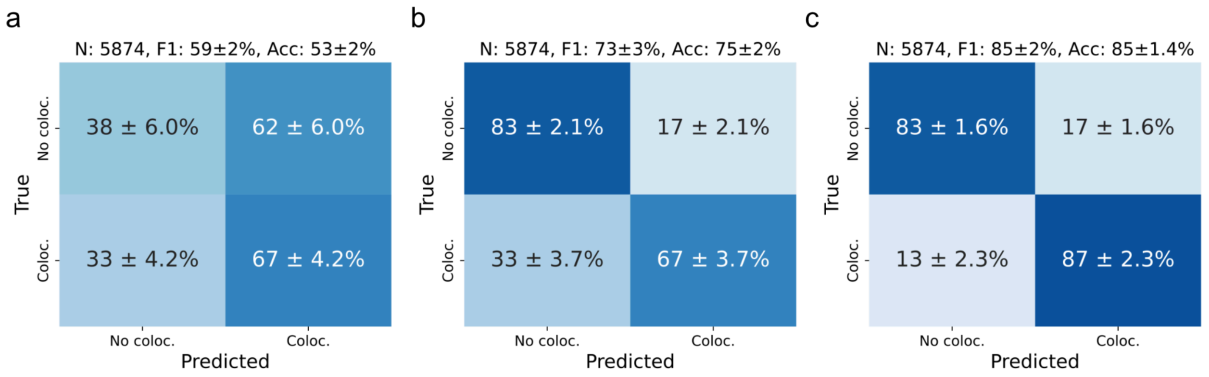


**Supplementary Figure 10:** Confusion matrices for the classification of true (Coloc.) and false (No coloc.) colocalization. **a)** Distance-based evaluation of classification using a random forest and average inter-trajectory distances. **b)** Pearson's correlation binarizing colocalization by accepting trajectories with correlation above 0.5 as true.(ref) **c)** Colocalization Fingerprinting. Additional classification metrics include accuracy (Acc), number of tracks (N), and F1-score (F1). Error bars depict standard deviations across a ten-fold cross-validation.

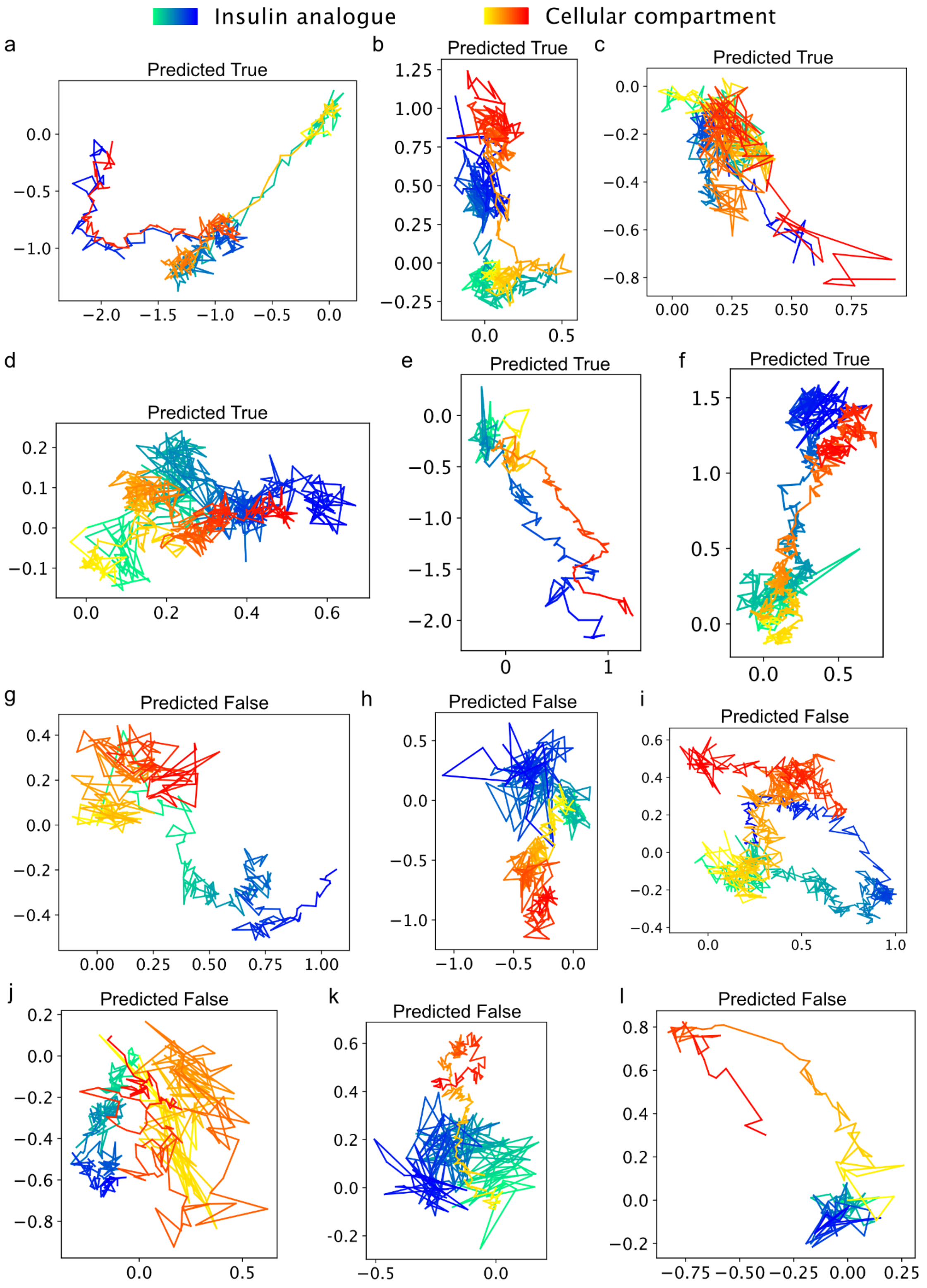


**Supplementary Figure 11:** Examples of insulin trajectory pairs evaluated by Colocalizational Fingerprinting either predicted true or false colocalization. Notably, trajectories with concerted motion are predicted as colocalizing, and trajectories with spatial proximity but independent motion are predicted as false colocalization. Trajectories are colored reflecting the temporal axis from light to darker shades. Blue shades are insulin trajectories and red shades are endosomal compartments. All coordinates are in micrometers.

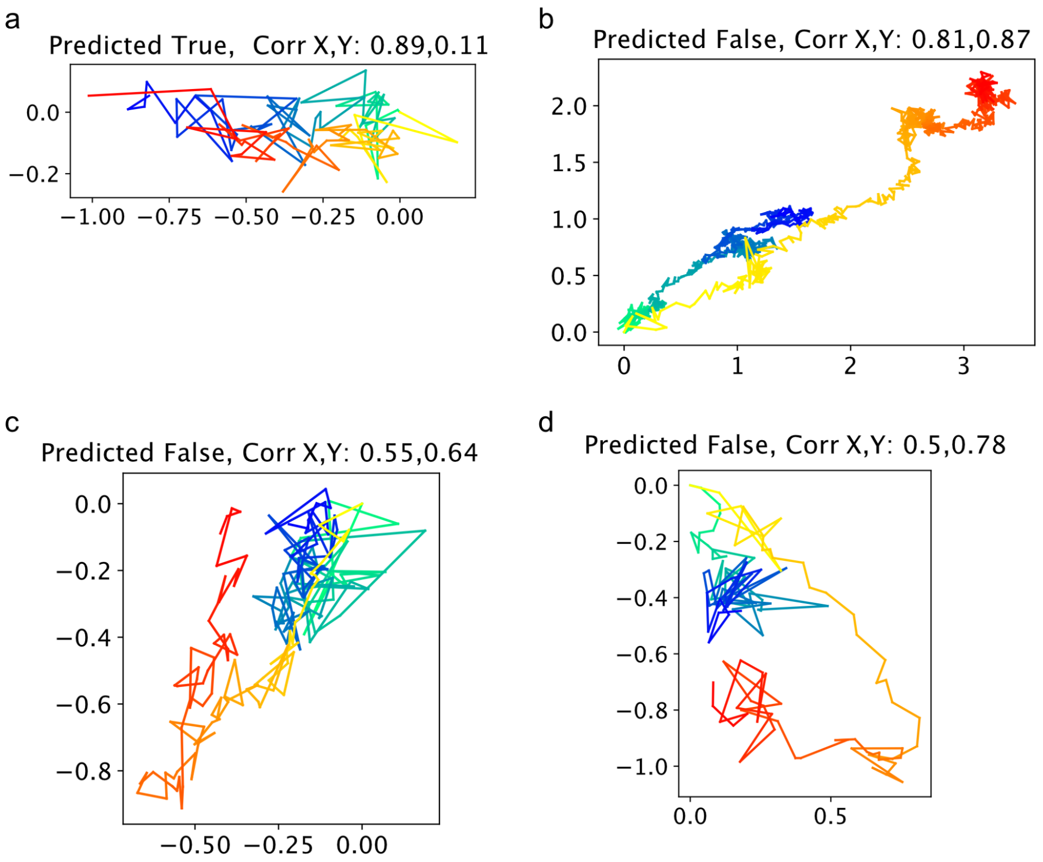


**Supplementary Figure 12:** Examples of insulin trajectory pairs evaluated by Colocalizational Fingerprinting either predicted true or false colocalization. Pearson's correlation (Corr X, Y) is included for the x and y coordinates, respectively. Notably, trajectories with low Pearson's correlation can exhibit concerted motion, yet obscured by noise (**a**). Likewise, trajectories with high Pearson's correlation can exhibit distinct, independent motion (**b-d)**. Combined, this highlights that Pearson's correlation can have flaws for experimental data convolved by crowding and noise. In contrast, Colocalization Fingerprinting accurately differentiates concerted from distinct motion (**a-d)**. Trajectories are colored reflecting the temporal axis from light to darker shades. Blue shades are insulin trajectories and red shades are endosomal compartments. All coordinates are in micrometers.

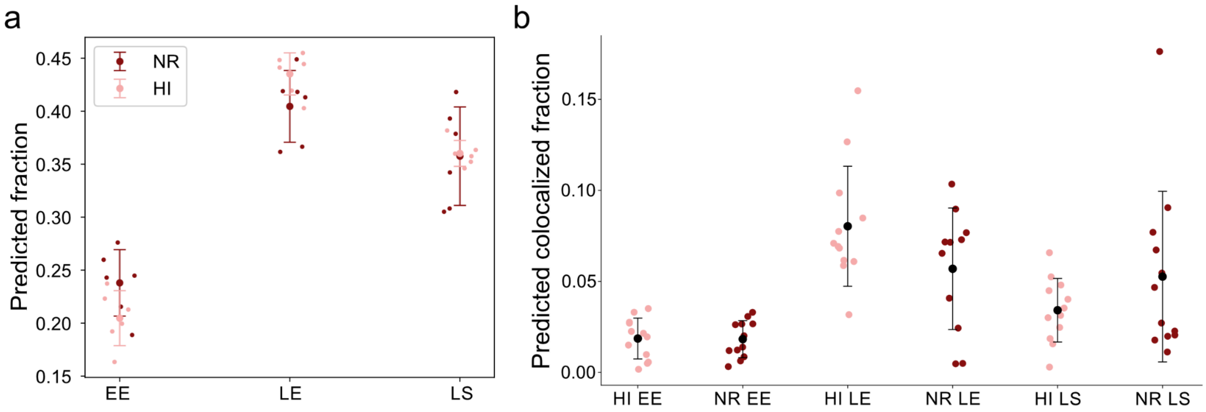


**Supplementary Figure 13:** Results for intracellular trafficking of insulin isoforms in HeLa cells. **a,** Fraction of tracks of either insulin analog, predicted by DeepSPT to diffuse like the respective compartment for trajectories above 50 time points. **b,** Colocalization propensities with endosomal compartments for the insulin analogs in HeLa cells. **a-b,** HI = human insulin, NR = insulin aspart/NovoRapid, EE = Early endosomes, LE = Late endosomes, LS = Lysosomes. Each point represents a technical replicate. All errors depict standard deviation across reported technical replicates.
